# Novel algal modular LOV domain proteins expand opto-biotechnological avenues for controlling of eukaryotic riboswitching, translational and proteolytic processes

**DOI:** 10.1101/2025.06.20.660589

**Authors:** Manisha, Rajani Singh, Suneel Kateriya

## Abstract

Light, Oxygen, or Voltage (LOV) domains mediate blue light-gated signal transduction, regulating diverse opto-biological functions. LOV domain functions either standalone or fused with effector domains, regulating the downstream signalling process. The current repertoire of LOV domain-based tools is limited to a relatively small number of naturally occurring proteins. In this study, we have identified novel algal LOV domain fused with different effector domains as potential light-mediated translational, ribogenetic and proteolytic switches, highlighting their unexplored avenues of opto-biotechnology. LOV-domain fusion with eIF4E suggests its potential as a light-controlled translational switch and as an opto-ribogenetic regulator. LOV-SppA might be used as a light-gated proteolytic switch. Additionally, LOV coupled with UFD1, UbiH, mannosyl-oligosaccharide glucosidase and biosynthetic gene cluster (BCG) molecular chassis pave the way for opto-biomanufacturing strategies of relevant valuable algal bioactive. Here, we report the discovery of 13 novel algal modular LOV domain-containing proteins across the algal system through comprehensive bioinformatics, bio-curation and systems biology approaches. It offers important insights into the structural and functional diversity of LOV photoreceptors in algae. Hence, these newly identified modular LOV domain-containing proteins expand the platform of opto-biotechnology applications. These finding lay the foundation for future research on the mechanistic basis for light-driven signalling cascade of RNA, translational and protein homeostasis in algae, and potentiate development of next-generation opto-biotechnological tools for synthetic biology, optogenetics and opto-biomanufacturing of valuable bioactive via regulation of biosynthetic gene cluster (BGC) in green lineage.

## 1 Introduction

Light has emerged as an exceptionally powerful tool in the fields of optogenetics, synthetic biology and opto-biotechnology due to its non-invasive nature (Tischer & Weiner, 2014a) . It imparts a high degree of spatial and temporal precision with the help of photoreceptors. Photoreceptors are light-sensitive proteins that absorbs different wavelength of light-mediating different signal transduction pathways. The photoreceptors naturally found in nature includes-BLUF (Blue Light sensors Using FAD), cryptochromes, rhodopsins, UVR8, phytochromes, and Light, Oxygen, or Voltage (LOV) domain-containing proteins (Briggs & Huala, 1999; Christie et al., 1998; Ebrey, n.d.; Hegemann et al., 2001). Among these, LOV domain-containing proteins, members of the Per-ARNT-Sim (PAS) domain superfamily, are blue light-sensitive proteins. It has garnered significant attention due to its small size (Shcherbakova et al., 2015), reversible and versatile photocycle (Buckley et al., 2015; Chapman et al., 2005). This photocycle varies from seconds to hours depending on the protein and environmental conditions (Zayner & Sosnick, 2014a; Zoltowski & Crane, 2008). This domain utilizes an endogenous, non-covalently bound flavin mononucleotide (FMN), flavin adenine dinucleotide (FAD), or riboflavin (RF) as chromophore (Mathony & Niopek, 2021; Tischer & Weiner, 2014b). Upon light absorption, the LOV molecules bind with the chromophore and undergo conformational changes, thereby modulating downstream biological activity (Kopka et al., 2017). These protein shows built-in modularity and are known to be distributed across three kingdoms of life (Krauss et al., 2009), showing a conserved GxNCRFLQ motif (Christie et al., 2002). Interestingly, LOV domain proteins are known to function as standalone or as multidomain proteins where they are coupled with a range of effector domains (Heintz & Schlichting, 2016). In multidomain form, LOV acts as a reversible sensory module and can be found located at N or C terminus (Takahashi et al., 2007; Vogt & Schippers, 2015) associated with various effector domains such as kinases (Christie, 2007; Djouani-Tahri et al., 2011), sulfate transporter/anti-sigma factor antagonists (Chan et al., 2013), <u>phosphodiesterases</u> (Tarutina et al., 2006) and phosphatases (Losi, 2013), and DNA-binding domains (HTH and HLH) (Motta-Mena et al., 2014), σ factor regulators (Ávila-Pérez et al., 2006) and Zinc fingers (Brunner & Káldi, 2008), F-box domains (Feke et al., 2021; Ito et al., 2012), proteins regulating circadian rhythms (Heintzen et al., 2001; Vaidya et al., 2011), cyclases (Raffelberg et al., 2013; Vide et al., 2023), transcription factors (Gligorovski et al., 2023; Herrou & Crosson, 2011) Thanks to its adaptable behaviour, LOV is a powerful tool in the synthetic biology and optogenetics fields. LOV-based domains are used in gene regulation (Wu et al., 2009), reversible control of protein-protein interactions (Shao, 2019) and protein localisation in live cells (Kennedy et al., 2010), light-activated switches in synthetic biological constructs (Lan et al., 2022), biosensors (Potzkei et al., 2012), photoactivatable enzymes (Lungu et al., 2012), and control of biofilm production (Moser et al., 2019). In addition, they are known to control cell physiology, reprogram metabolism (Zhao et al., 2018), promoting cell growth and tissue patterning (Krueger et al., 2019) and control of cell ablation. (Buckley et al., 2017; Li et al., 2021).

Despite their widespread use, the current repertoire of LOV domain-based tools is limited to a relatively small number of naturally occurring proteins, many of which were not originally evolved for optogenetic functions. Consequently, achieving optimal dynamic range, specificity, and modularity in engineered systems remains challenging. Additionally, most existing LOV domains are constrained by their natural domain architecture and are not easily adaptable to synthetic modular designs. In this context, the discovery and characterization of novel modular LOV domain proteins, those that exhibit diverse architectures, functional domains, and regulatory capabilities, represent a promising avenue for expanding the optogenetic toolbox. Modular proteins with customizable or naturally diverse domain arrangements could offer integration of synthetic circuits with fine-tuned spatiotemporal control. Thus, understanding the variety of effector functions and domain arrangements will reveal how cells adapt to light and how blue-light signals are transduced through modular architectures.

Here, we report the identification and molecular characterization of uncharacterized modular LOV domain-containing proteins by deploying an array of bioinformatics, bio-curation and systems biology approaches. Using a combination of database mining, homology analysis, domain architecture characterization, comparative structural insight and interactome with biosynthetic gene cluster components, we uncover their potential for next-generation opto-biotechnological tools. Further, with the help of bio-curation of protein networking, we show their broader application in synthetic biology including biomanufacturing of bioactive simply by illumination. Our findings lay the groundwork for designing customizable, light-controlled systems with broader applicability in synthetic biology and opto-biotechnology area.

## 2. Methodology

### 2.1 Identification of novel algal modular LOV domain proteins

To identify the novel algal modular LOV domain containing proteins, the uncharacterised LOV-domain containing protein sequences were retrieved from the Joint Genome Institute (JGI) PhycoCosm portal (https://phycocosm.jgi.doe.gov) and National Center for Biotechnology Information (NCBI) protein database (https://www.ncbi.nlm.nih.gov/protein/) using *Chlamydomonas reinhardtii* LOV protein as a query sequence. Further, the sequences were subjected to domain prediction using the Conserved Domain Database (CDD) via the CDART (Conserved Domain Architecture Retrieval Tool) (https://www.ncbi.nlm.nih.gov/Structure/lexington/lexington.cgi) to check the authenticity of LOV proteins coupled with diverse effector domains.

### 2.2 Homology analysis and motif discovery of algal modular LOV domain

Homology between the LOV domains for each protein was analysed through multiple sequence alignment using CLUSTAL W in BioEdit tool (Dagona, 1999). The conserved motifs across LOV domain sequences were identified using Multiple Expectation maximization for Motif Elicitation (MEME) suite (Version 5.5.8) (Bailey et al., 2009). With the zero or one occurrence per sequence (zoops) model, parameters were set to detect up to three motifs with a width of 6–50 amino acids. Only highly conserved motifs (E<1e-5) were taken into consideration for analysis, and the significance of motifs was assessed using E-values.

### 2.3 Comparative structural analysis of algal modular LOV domain proteins

The LOV domain sequences of different proteins were submitted to AlphaFold server for the generation of 3-dimensional models (https://alphafoldserver.com/). Structural superimposition and visualisation were performed using PyMOL (https://pymol.org/). Structural conservation and alignment of key secondary structural elements (α-helices and β-sheets) were analysed to compare core architecture across diverse LOV proteins.

### 2.4 Evolutionary analysis of algal modular LOV domain-containing protein

Phylogenetic analysis was done for LOV domain and effector domains of modular proteins using MEGA 12 (Kumar et al., 2024). Alignment of different domains was done in CLUSTAL W tool in MEGA12 and further analysis was performed by employing the Maximum Likelihood method, based on the JTT matrix-based model (Jones et al., 1992) with 1000 replicates. The JTT matrix of pairwise distance was subjected to Neighbour-Join and BioNJ algorithms for the construction of the first tree(s), which was then used to select the topology with a higher log likelihood value.

### 2.5 Prediction of protein-protein interaction for the proposed functional model based on coupled effector domain

Sequences retrieved from JGI and NCBI of different effector domain were submitted to string 11 software for protein-protein interaction analysis (PPI) (STRING v11: protein-protein association networks with increased coverage, supporting functional discovery in genome-wide experimental datasets. (Szklarczyk et al., 2019). Further, network visualization was done using Cytoscape V 3.10.3 (Shannon et al., 2003).

### 2.6 Bio-curation of crosstalk between LOV-coupled effector domains with biosynthetic gene clusters mediated metabolic pathways

The crosstalk between LOV-coupled effector domain and BGCs was demonstrated in *Chlamydomonas reinhardtii* by creating a bio-curated protein network prediction using String database version 12.0 (https://string-db.org/). For protein network prediction, the sequence of selected LOV-coupled effector domains, important molecules/enzymes of the selected metabolic pathway and core domain proteins of BGCs identified by (Singh et al., 2025) were retrieved from the JGI genome and the NCBI database and submitted to the String database. The obtained network was analysed and refined through Cytoscape (Version 3.10.3). The betweenness centrality metric with default layout and shortest path was employed to identify the top 20 key nodes bridging the network. The scores ranked for interacting partners using the betweenness method are indicated in colour, ranging from red to yellow.

## 3. Results and Discussion

### 3.1 Novel algal LOV modules are coupled with diverse effector domains

In the present study, we have identified novel algal modular LOV domain-containing proteins, coupled with diverse effector domains. A total of 13 uncharacterized modular LOV proteins were identified. These identified effectors are involved in redox homeostasis, protein regulation, ubiquitin-mediated degradation, energy metabolism, cell cycle regulation, transportation, RNA-binding motifs, stress adaptation, etc. The detailed information about their structure (domain architecture), source organism, function and accession number is listed in the Table 1. Their homology and applications are discussed in detail in the further relevant sections.

**Table 1.**
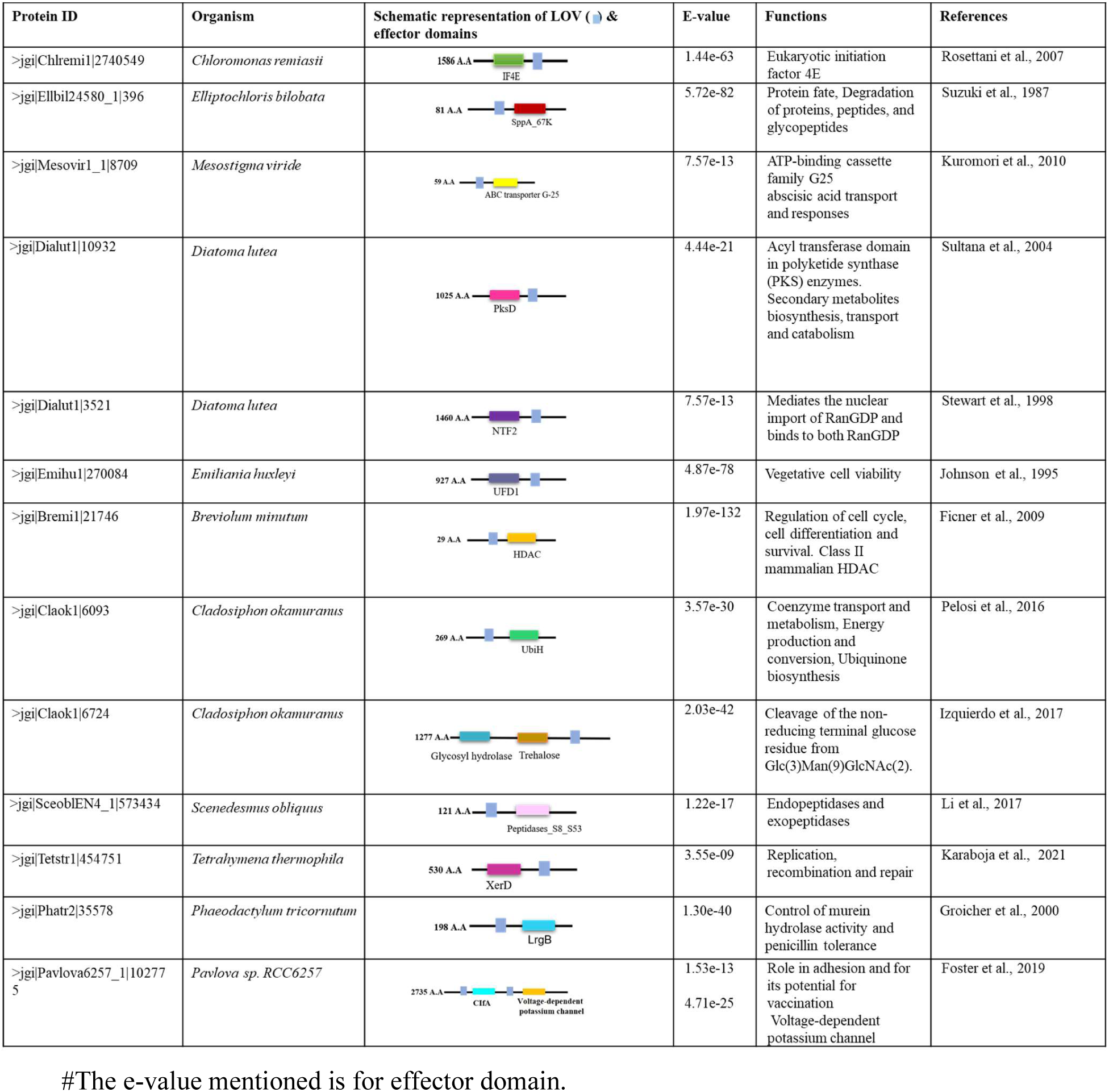
List of identified algal modular LOV containing domains consisting of their function, schematic structural representation and source organism with their accession ID.

### 3.2 Molecular homology and variability analysis provide novel insight into photodynamic properties of the different modular LOV proteins

To discover novel modular LOV-domain containing proteins coupled with different effector domains as potential optogenetic tools, homology analysis between the LOV domains of modular LOV proteins was performed. The LOV1 of *C. reinhardtii* was used as a query to mine the algal genome, consisting of different effector domains coupled with LOV. The analysis indicates that the amino acid residues essential for chromophore binding and photocycle is conserved throughout the identified modular LOV protein. The cysteine residue important for photoadduct formation is embedded within the NCRFLQ motif (Fig. 1). The conserved NCRFLQ region and surrounding residues (highlighted with star) are likely essential for cofactor binding (FMN) and proper domain folding. Apart from cysteine (marked with arrow), glutamine (Q) except in HDAC effector domain, histidine (H), and asparagine (N) residues were also conserved, suggesting their roles in light-sensing and signal transduction. This suggests that important residues for photoadduct formation and photocycle are conserved across the algal system, in spite of having varied effector domains. The variability in the function and signal transduction of LOV protein is due to domain shuffling and coupling with diverse effector domains. We further investigated the difference in the motif present in them. The result suggested that the motif for adduct formation (motif 1 and motif 2) is conserved (Fig. 1B). Our result corroborated with that of similar publication (Glantz et al., 2016). They reported that motifs related to adduct formation and photocycling (motifs 1 and 2) are conserved among the 18 well-characterised LOV domains. The mutation in these domains leads to impairment of adduct formation and photocycle (Glantz et al., 2016). However, motif 2 was absent in UFD1 coupled LOV domain. This suggests that this modular LOV protein can be used for altering the downstream signalling or biological process as per the cell’s requirement. Notably, we identified a new motif (motif 3) in all the LOV domains coupled with different effector domains except LOV-XerD. It would be interesting to see how this motif affects the overall light sensing and transduction mechanism in an organism.

**Fig. 1:**
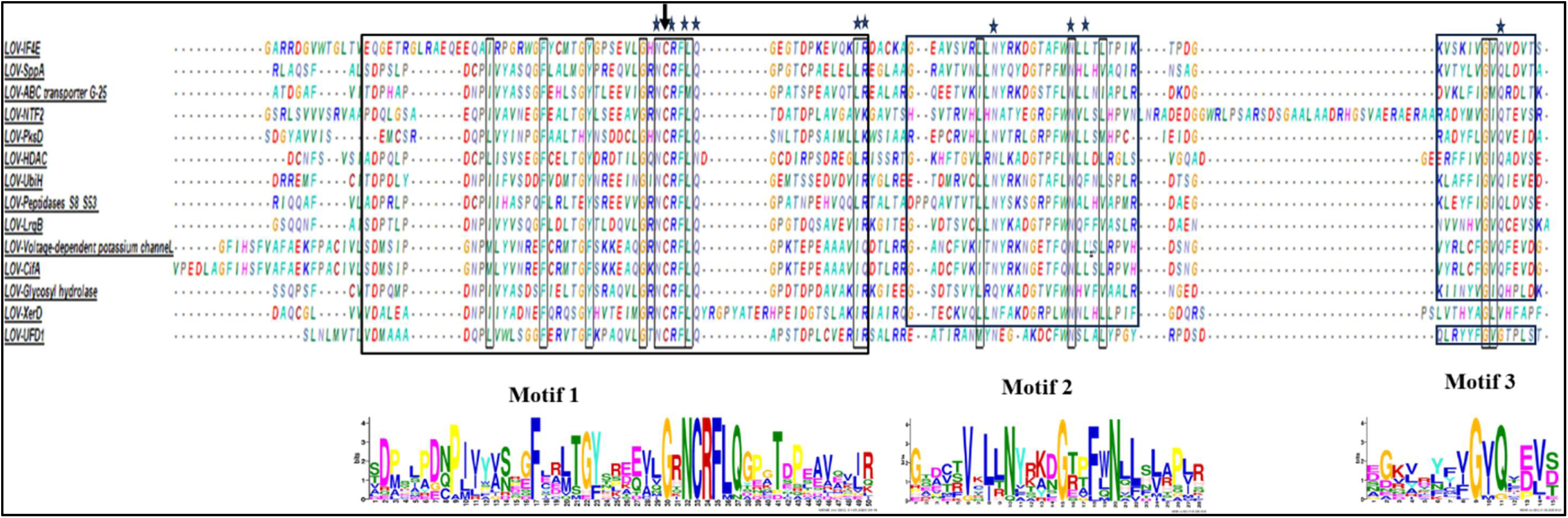
Homology analysis of different LOV domains of modular proteins using Clustal W in BioEdit software. Residues important for chromophore binding and photocycle are marked with star. The black arrowhead indicates a conserved cysteine residue interacting with FMN present in all the LOV1 sequences. Motif Elucidation using MEME software showed the presence of three conserved motifs marked with black boxes, and sequence logos for each motif are shown below in the picture.

### 3.3 Structural analysis of novel algal LOV domain suggests diverse photo-chemical characteristics and their putative function

The core structure of LOV domain is conserved throughout all the domains. It comprises 4 alpha helices surrounded by 5 central beta anti-parallel sheets and has a flavin binding pocket (FMN) (Hart & Gardner, 2021). In the present study, we conducted a comparative structural analysis of the LOV domain of all identified modular proteins. The superimposition of the LOV domain of all the selected modular proteins shows conserved alpha helices, anti-parallel beta sheets and FMN binding pocket (Fig. 2). However, variability was observed in the loop region. The loop region was found to be elongated at N-terminus in IF4E and voltage-dependent effector modular LOV-domain containing proteins. Moreover, the loop region of NTF2 at C-terminal was also larger than the others. The loop region enables functional versatility in the LOV domain. The residues of the loop emerging from the alpha and beta regions (E27 and I66) control the lifetime of the adduct form by modulation of hydrogen bonding in *Pseudomonas putida* (Arinkin et al., 2017, 2021). Similarly, Pro 426 residue of loop region maintains structural integrity at the time of dark-light transition state. Additionally, interaction of the LOV domain with the loop region acts as signal transduction for the downstream process (Arinkin et al., 2017). Thus, the loop region plays an important role in photocycle regulation. This suggests that the different modular LOV proteins with variable effector domains can provide a wide choice for optogenetic applications.

**Fig. 2:**
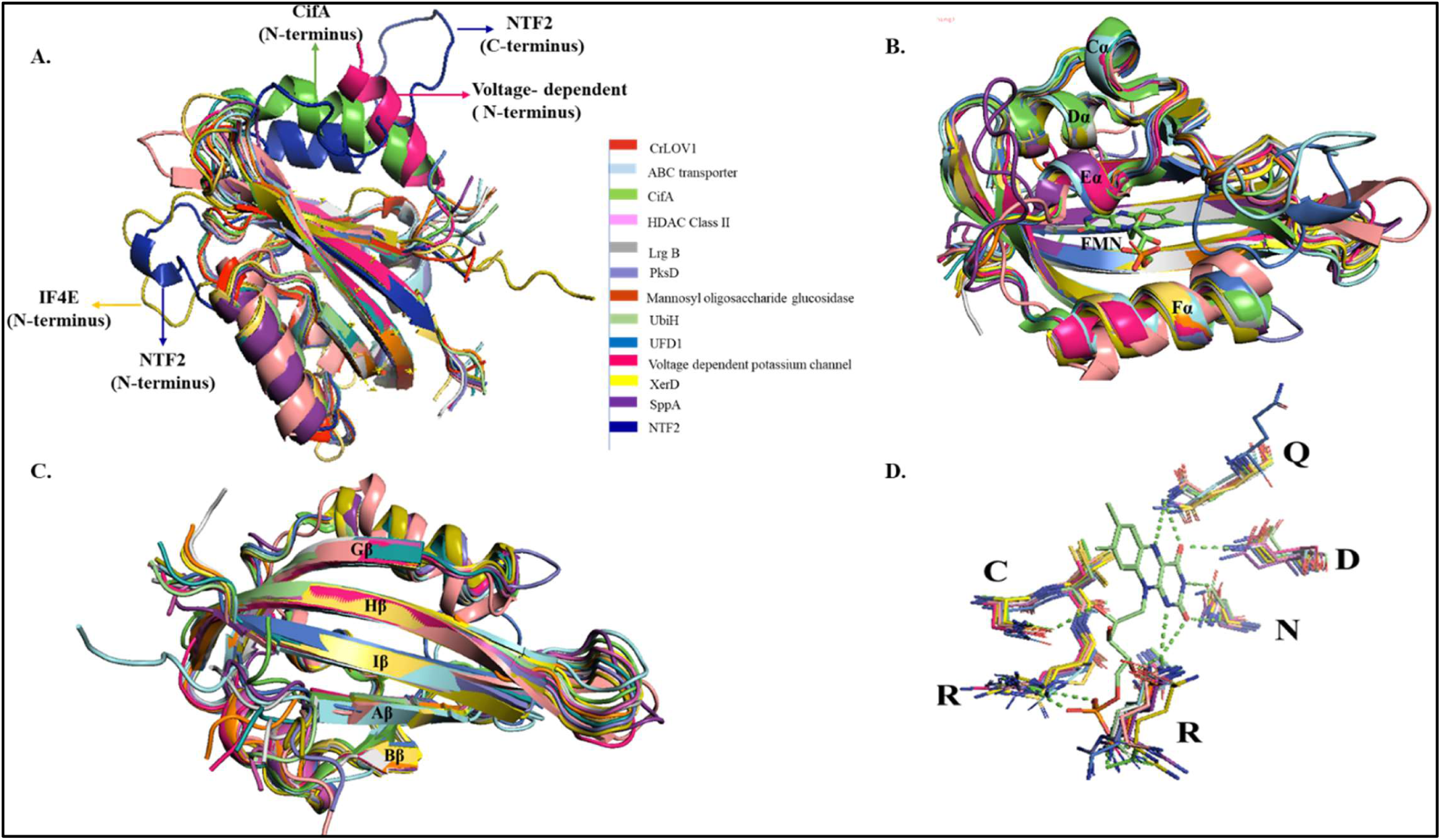
Structure prediction and their comparative analysis of LOV domain of different modular proteins. A. Superposition of all LOV domains. B. Superposition of LOV domains of all modular proteins showing conserved alpha helices and FMN (Flavin mononucleotide) binding pocket. C. Superposition of Flavin binding pocket showing residues corresponding to CrLOV1. D. Structure showing conserved antiparallel beta-sheets in all LOV domains with variable regions in loops. Variable regions are marked among different domains.

Apart from loop regions, variability was also observed for helix region of CifA, NTF2 and voltage-dependent potassium channel at their N-terminus. Helices are important for structural stability, signal transduction and interaction with beta sheets (Kim et al., 2024; Zayner et al., 2019). Variability in these regions alters the light-sensing ability and signal propagation to the effector domain. They are important for allosteric regulation of the effector domain (kinase in the case of phototropin) and signal integration. Thus, the dynamic behaviour of modular LOV domains underlies their light-dependent switching behaviour, making it a potent optogenetic tool (Kim et al., 2024; Zayner et al., 2019).

Further, the superimposition of FMN pocket binding shows that residues important for chromophore binding and photocycle show correspondence to CrLOV domain. The comparative results for the residues are shown in Table S1. The R63, Q120, and N89 are crucial for chromophore finding and photocycle varies in UFD1 (Q120-G), mannosyl-oligosaccharide glucosidase (N89-Q), XerD (Q120-V) and CifA and voltage-dependent potassium channel (R63Q). Asn, Arg and Gln are important for hydrogen bond formation with FMN (Fedorov et al., 2003; Raffelberg et al., 2011; Zayner & Sosnick, 2014b). They are highly conserved across LOV domains. The mutation in these residues alters the chromophore binding and dark-state recovery. For example, variation in residue from N89Q in mannosyl-oligosaccharide glucosidase might result in accelerated dark-state recovery. Further research is needed in this direction. Hence, our data unveil a novel modular LOV domain-containing protein for new optogenetic tool development.

### 3.4 Evolutionary insight indicates origin of novel modularity of algal LOV domain and effector domains

To explore the evolutionary trajectory of modular LOV domain-containing proteins, we constructed a phylogenetic tree based on their LOV domain sequence. The tree consists of a total of five clades (Fig. 3). Among them, Clade I, LOV-SppA, LOV-Peptidases and LOV-ABC transporter are observed as a prominent clade. They are involved in light-regulated proteolytic cleavage and transportation. Clade II comprises of LOV-mannosyl oligosaccharide glucosidase, HDAC (histone deacetylase) and LrgB. Further, LOV-voltage potassium channel and LOV-CifA (Cytoplasmic incompatibility factor A) are different effector domains belonging to the same algae, clade with UbiH (Ubiquinone flavin monooxygenases) (Clade III). Similary, LOV-NTF2 and LOV-PksD belong to the same algae, i.e, *Diatoma lutea* (occurs both in fresh water and marine) clades with LOV-XerD from *Phaetodactylum tricornatum* (marine) and LOV-UFD1 from *Emiliania huxleyi*. Interestingly, LOV-IF4E forms another individual clade (Clade IV), suggesting an independently evolved light-dependent mechanism of translational regulation, specific to green algal lineages. The LOV-UbiH and LOV-mannosyl oligosaccharide glucosidase both belongs to *Cladosiphon okamuranus* (Phaeophyceae, brown algae) cluster in the same lineage, suggesting a shared functional theme in redox and glycan processing regulated by light within this phylum. Meanwhile, LOV-HDAC from *Breviolum minutum* (Dinoflagellata) and LOV-UFD1 from *Emiliania huxleyi* (Haptophyta) suggest the evolution of LOV fusions involved in chromatin remodelling and protein degradation, respectively. Notably, LOV-CifA and LOV-voltage-gated potassium channel, despite having distinct effector domains, both derive from *Pavlova* sp. RCC6257 (Haptophyta), highlighting domain modularity within the same phylum. This implies possible genomic co-localisation and diversification of LOV-based signalling components in response to light. The phylogenetic tree for HDAC and peptidase S8 is given in Fig. S1. The protein accession number and source organism used for phylogenetic analysis and homology analysis of different effector domains are given in supplementary Table S2. Overall, the evolutionary tree suggests LOV domain as modular hub coupled with distinct effector domain across different lineage to satisfy the diverse physiological need of organisms.

**Fig. 3:**
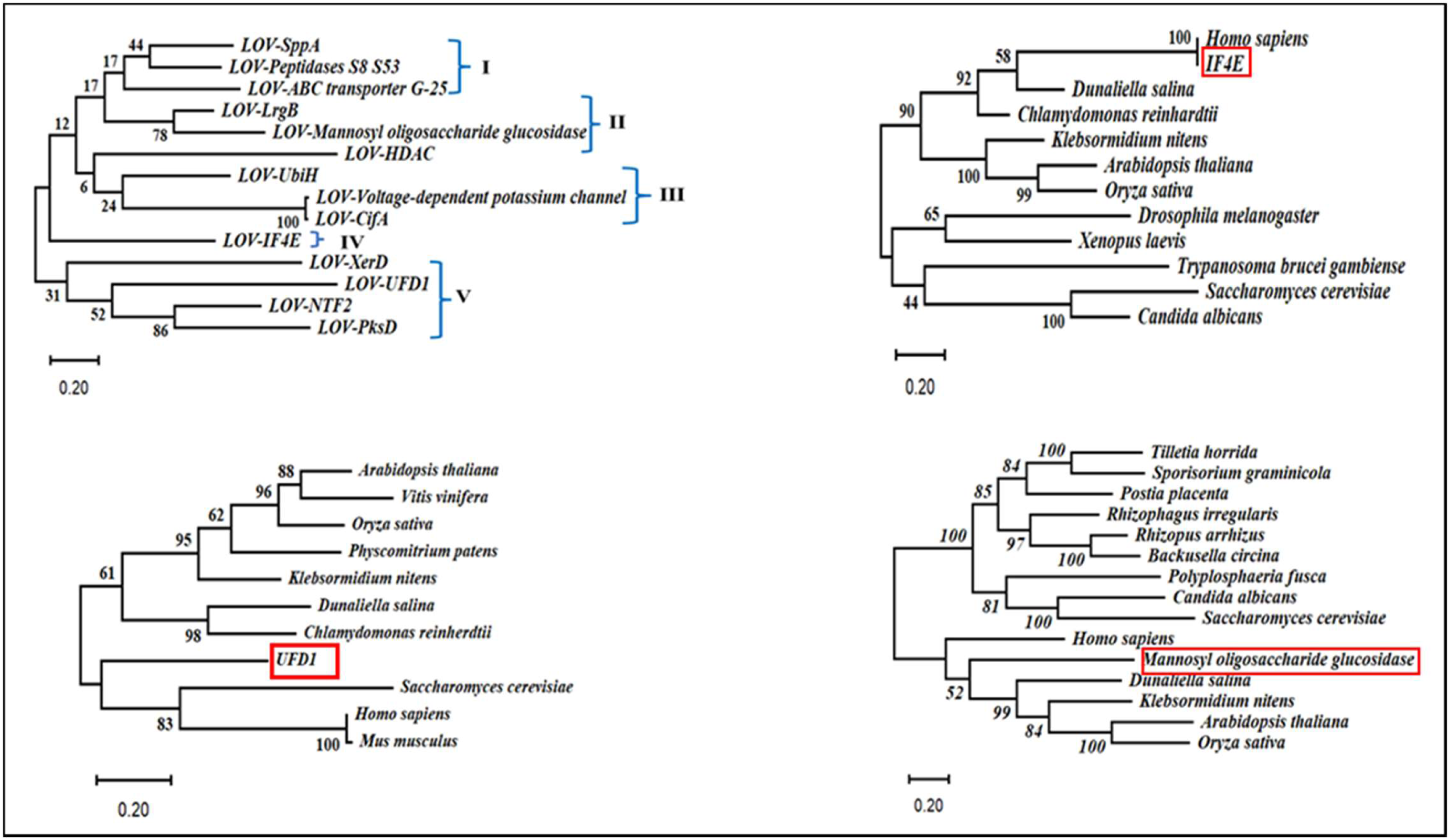
Phylogenetic analysis of LOV domain of all modular proteins. Alignment of protein sequences was carried out using ClustalW in MEGA12. The tree was analyzed using Maximum likelihood method and Jonas –Taylor-Thorton model with 1000 bootstrap replicates. Evolutionary analyses were conducted in using MEGA 12 software.

### 3.5 Homology analysis of effector domain reveals its implication as unexplored optogenetic potential

Homology analysis was done to predict the functional compatibility, potential for engineering and their evolutionary significance. We identified that the functional residues, such as catalytic residues and motifs, are conserved across organisms (Fig. 4 and S2-S4). The effector domain IF4E, eukaryotic initiation factor, regulates protein synthesis. It is extensively studied in yeast and mammals to understand its ability to modulate protein synthesis. To initiate translation, it binds with the m7GpppN cap at the mRNA. The homology analysis suggests that amino acids involved in m^7^GpppN binding for stabilizing the structure of the protein are conserved across the organisms (Fig. 4A). These are either involved in tertiary interaction or located in the hydrophobic core (Monzingo et al., 2007). Similarly, mannosyl oligosaccharide glucosidase homology reveals that residues important for α-glucosidase activity are conserved in yeast, fungi, algae, higher plants and humans (Fig. 4B). It is involved in the glycan processing. MOGS is highly specific to its oligosaccharide substrate. Hence, this effector domain can be utilized for targeted light-responsive glycosidase. Furthermore, in UbiH, Ubiquinone flavin monooxygenases, the DG amino acid sequence (play dual function in FAD and NAD(P)H binding), the GXGXXG motif of well-known Rossmann fold, GD sequence for FAD binding motif having highly conserved aspartyl residue that contacts the O-3′ of the ribose moiety of FAD found to be conserved among different organisms (Fig. S2). UbiH is a crucial molecule involved in cellular bioenergetics, maintaining electrochemical gradient in respiratory chains from prokaryotes to eukaryotes (Gin et al., 2003; van Berkel et al., 2006). In our study, the important amino acids are conserved from *E. coli* to freshwater alga (*C. reinhardtii*), terrestrial alga (*K. nitens*) and higher plants (*O. sativa*). Thus, it shows wide application for various organisms. Likewise, another effector domain, UFD1, Ubiquitin fusion degradation protein 1 (UFD1), an important recognition component of proteolytic degradation, also has conserved homology for psi β-barrel motif and the α/βroll motif, which has binding sites for poly-ubiquitin chains (Fig. S3). Our result also corroborates with (Weihofen et al., 2000). Similarly, the homology analysis of the SppA effector domain (signal peptide peptidase) shows that the domain framework was conserved (Fig. S4). Thus, this shows that the identified novel effector domains can work efficiently as the amino acid sequence, residues and domain architecture are conserved. So when fused with the LOV domain, it might emerge as a potential optogenetic tool regulating diverse physiological functions.

**Fig. 4A:**
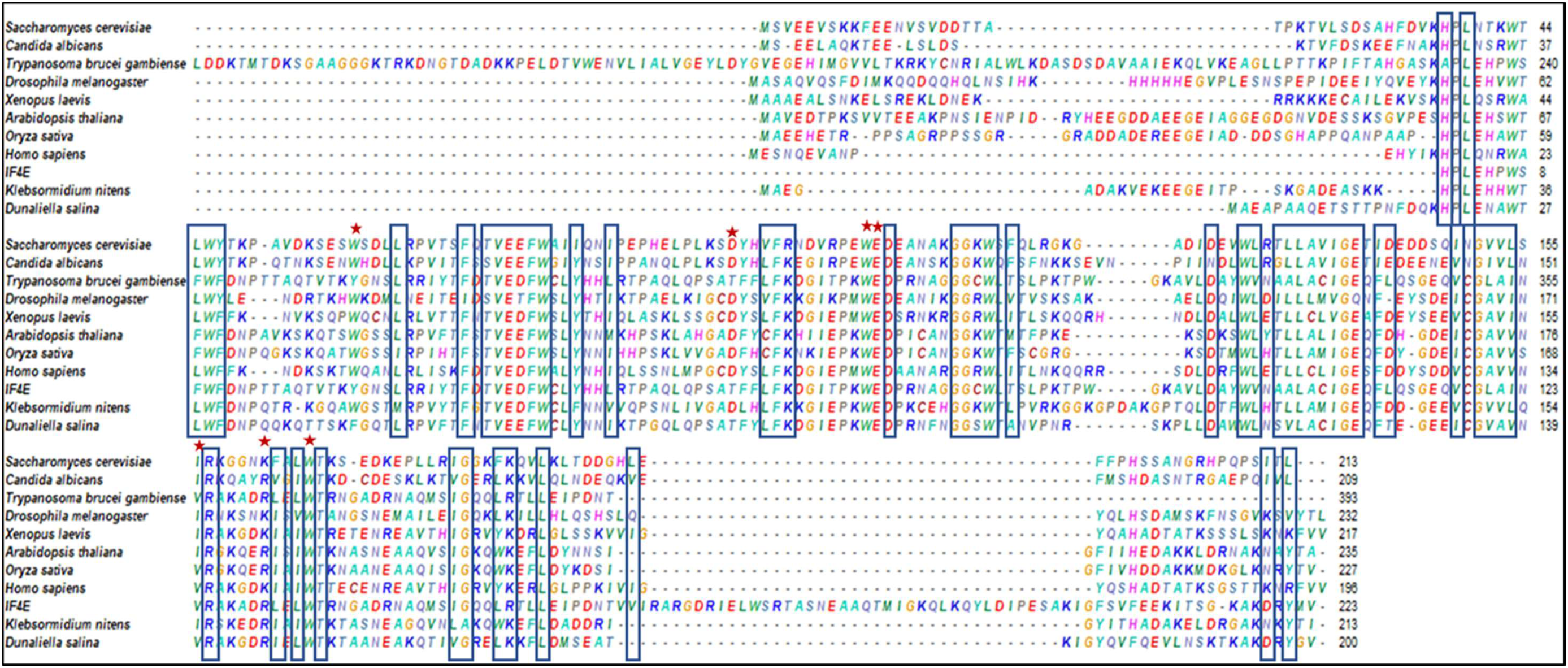
Homology analysis of different effector domain IF4E. Amino acids that are most directly involved in binding m^7^GTP shown through star. Amino acids that are important for stabilizing the structure of the protein are boxed in black; these are either located in the hydrophobic core or involved in other tertiary interactions in black box.

**Fig. 4B:**
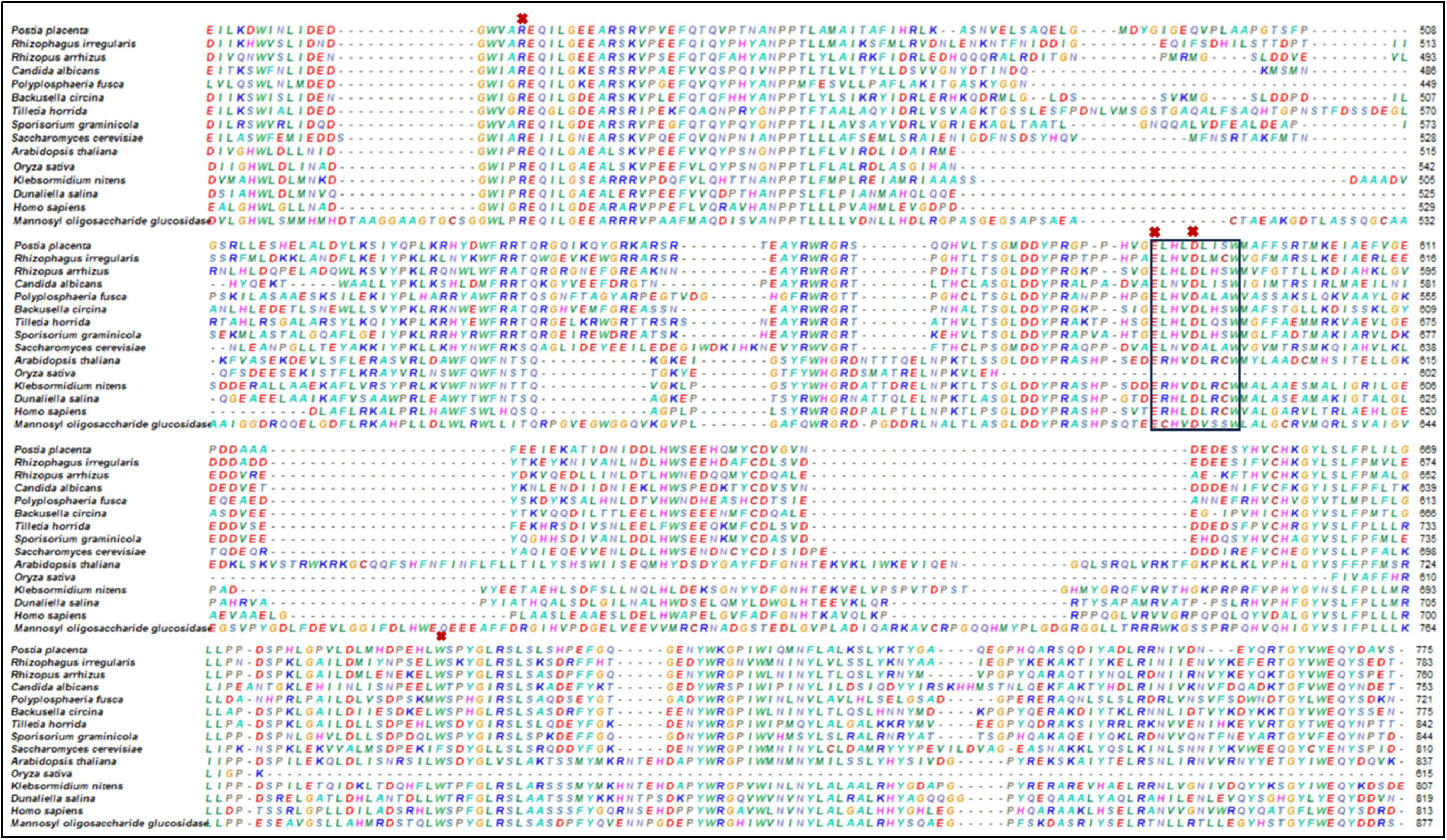
Shows homology analysis of mannosyl oligosaccharide glucosidase effector domain. Here, the black box represents the motif involved in substrate binding. Cross indicates conserved residues important for α-glucosidase activity.

### 3.6 Novel algal modular LOV-coupled effector domains: Unlocking potential for opto-biotechnology avenues

Coupling of the LOV domain with diverse effector domains offers a promising platform for light-inducible tools in synthetic biology, optogenetics, opto-biomanufacturing and opto-biotechnology. The identified novel modular LOV coupled with effector domains might enable, tuneable, reversible, targeted and precise control of diverse physiological processes. The integration of enzymatic, signalling and/or degradation-based effector domains with LOV paves the way for innovative opto-biotechnology applications in spatiotemporal control of cellular pathways such as protein turnover, metabolic engineering, etc. The predicted application of these domains are described below:

#### 3.6.1 LOV-eIF4E - light-controlled translation switch

Among the wide variety of modular LOV-containing proteins found during the present study, LOV-eIF4E is a particularly unusual combination of a eukaryotic translation initiation factor 4E (eIF4E) and a LOV (Light-Oxygen-Voltage) sensor domain. As per our knowledge, it is the first naturally occurring flavin-binding protein that is directly linked to a core translation regulator, signifying a special connection between mRNA translation and photoreception. This fusion increases the likelihood of light-regulated translational control through a LOV-mediated mechanism, as eIF4E plays a crucial role in binding the 5′ cap structure of mRNAs to start translation. The PAL (PAS-ANTAR-LOV) proteins from *Nostoc multipartita* exhibit a similar evolutionary and molecular pattern to our observation (Weber et al., 2019). In these proteins, the ANTAR domain mediates RNA-binding activity that is allosterically modulated by the LOV domain in response to light. Although it has been demonstrated that PAL proteins control gene expression at the RNA level, the LOV-eIF4E protein offers a radically different regulatory method from traditional light-responsive transcription factors or RNA-binding proteins by implying a direct coupling of light perception to cap-dependent translational initiation.

#### 3.6.2 Novel modular LOV proteins as potential tool for opto-ribogenetics

Opto-ribogenetics, nexus optogenetics and ribogenetics (Pilsl et al., 2020). Opto-ribogenetic systems use photoresponsive protein-RNA interactions to precisely control the spatiotemporal behaviour of RNA, including translation, splicing, and degradation (Liu et al., 2022). Usually, these engineered systems depend on interactions between LOV caused by artificial light. Synthetic light-induced interactions between LOV domains and RNA aptamers inserted into target transcripts are usually the basis for these engineered systems. Nonetheless, the identification of a spontaneous LOV-eIF4E fusion implies that nature has already developed its own opto-ribogenetic systems, allowing for light-gated, reversible regulation of mRNA translation.

The novel modular LOV protein module, LOV-eIF4E module, identified in our study might work mechanistically by causing conformational changes in the LOV domain that either identify, reveal or conceal the cap-binding pocket of eIF4E, or else alter its interaction with partners in translation initiation complexes (like eIF4G). The LOV-eIF4E fusion is a natural model for the creation of photo-controllable translational switches from the standpoint of synthetic biology (Andres et al., 2019). This architecture may be used to create chimeric LOV-eIF4E proteins with specific RNA-targeting and light-responsiveness, opening up possibilities for metabolic engineering, optogenetic circuit design, and programmable gene expression. Furthermore, its discovery opens the door for further research into light-gated post-transcriptional regulation in various eukaryotic systems and broadens the scope of LOV domain functional diversity.

#### 3.6.3 LOV-signal peptide peptidase (SppA) shows plausible use as light-controlled proteolytic switches

SppA is signal peptide peptidase required for regulated intramembrane proteolysis (RIP). It cleaves the signal peptides after proteolytic removal from the exported protein. It occurs in algae, fungi, plants, animals and protozoa (Voss et al., 2013; Weihofen et al., 2000). In the fungus *Aspergillus nidulans*, it is an aspartyl protease that is crucial for mechanism of hypoxial adaptation and cleavage of sterol regulatory element binding protein (Bat-Ochir et al., 2016). In *E. coli*, it acts as serine protease (Wang et al., 2008). In *Arabidopsis thaliana*, it is identified as a light-regulated chloroplast protease (Lensch et al., 2001). By coupling of SppA with LOV, the protease activity can be controlled by switching on or off blue light. Thus, it can control targeted protein degradation, which is a crucial regulatory phenomenon in various cellular processes, controlling the quality and quantity of protein, specifically under stress conditions. Additionally, the spatial-temporal aspects of protein degradation can dissect the role of protein turnover in a particular cellular process. Thus, it can broaden our understanding of stress responsiveness, signalling and adaptation mechanisms in various organisms.

#### 3.6.4 LOV-domain as an opto-biotechnological hub for regulating glycan, fatty acid metabolism and energy metabolism in green lineage

The present work renders a LOV-mediated next-generation opto-biotechnology strategy to modulate the activity of Biosynthetic Gene Clusters (BGCs) responsible for regulating glycan, lipid, and energy metabolism in *C. reinhardtii.* Biosynthetic gene clusters (BGCs) collectively encode the genes and the enzymes for the biosynthetic pathway of a specific metabolite (Singh et al., 2025) have identified 2 BGCs in *C. reinhardtii* and shown their interaction with the photoreceptors with the help of curated PPI. In this study, we have shown light-regulated BGCs mediated control of effector domains, mannosyl oligosaccharide glucosidase (glycosyl hydrolase family 63 and trehalase), UFD1, and UbiH via bio-curated PPI using the String database and analysed it in Cytoscape to identify nodes regulating the overall network. (Fig. 5). The bio-curated PPI networking shows that alpha-1,4 glucan phosphorylase (PBOS), an allosteric enzyme of carbohydrate metabolism, interacts with SQD2, which contains the BGC core domain glycosyl-trans 1-4 of saccharide production. Also, it shows interaction with the photosynthesis pathway. Interestingly, the BGC core domain fatty acid desaturase shows interaction with phytoene synthase, which is directly regulated by phototropin. The UbiH effector domain interacting partner CAS1, which is a terpene cyclase/mutase family member, shows direct interaction with carotenoid metabolism (zeaxanthin epoxidase, ZEP and phytoene synthase, PSY). Additionally, the members of UFD1, RPL40, a ribosomal binding protein, directly interact with Cop4 photoreceptor, ERF1 of BGC core domain protein and lycopene ε-cyclase (LYCE) of carotenoid metabolism. UFD1 and UbiH are involved in the proteosomal degradation of protein. They are important components of ubiquitin-mediated proteolytic cleavage and are involved in numerous biosynthetic pathways. Therefore, it is hypothesised that the LOV domain can also be applied as an optogenetic tool for regulating glycan processing and fatty acid metabolism, in turn, their production in other microalgae and the green lineage system. Similarly, the bio-curated PPI for another BGC cluster identified in *C. reinhardtii* also shows their interaction with different photoreceptors such as phototropin, Chlamyopsin 4 and UVRB (Fig. 6). Interestingly, they are linked with different core domain components identified for putative cluster. Additionally, we have also shown the interacting partners of each effector domain involved in the bio-curated BGCs-mediated networking (Fig. S5). The interacting partners of these effector domains show their involvement in diverse cellular processes. Therefore, the integration of LOV-domain-based photoreceptors with core metabolic enzymes such as fatty acyl desaturase and mannosyl oligosaccharide glucosidase, the latter bearing glycosyl hydrolase and trehalase domains, enables dynamic, light-tunable control over essential biosynthetic functions. The inclusion of UFD1 and UbiH, key effectors in ubiquitin-mediated degradation pathways, further adds a layer of reversible post-translational regulation, allowing selective degradation or stabilization of pathway components in response to illumination. This modular framework not only provides spatiotemporal precision over metabolic outputs but also aligns with the architecture of BGCs, making it highly adaptable for rewiring natural product biosynthesis and metabolic flux. Collectively, this design establishes a powerful platform for light-responsive, non-genetic control of metabolism, paving the way for customizable applications in synthetic biology, precision bioengineering, and sustainable biomanufacturing.

**Fig. 5:**
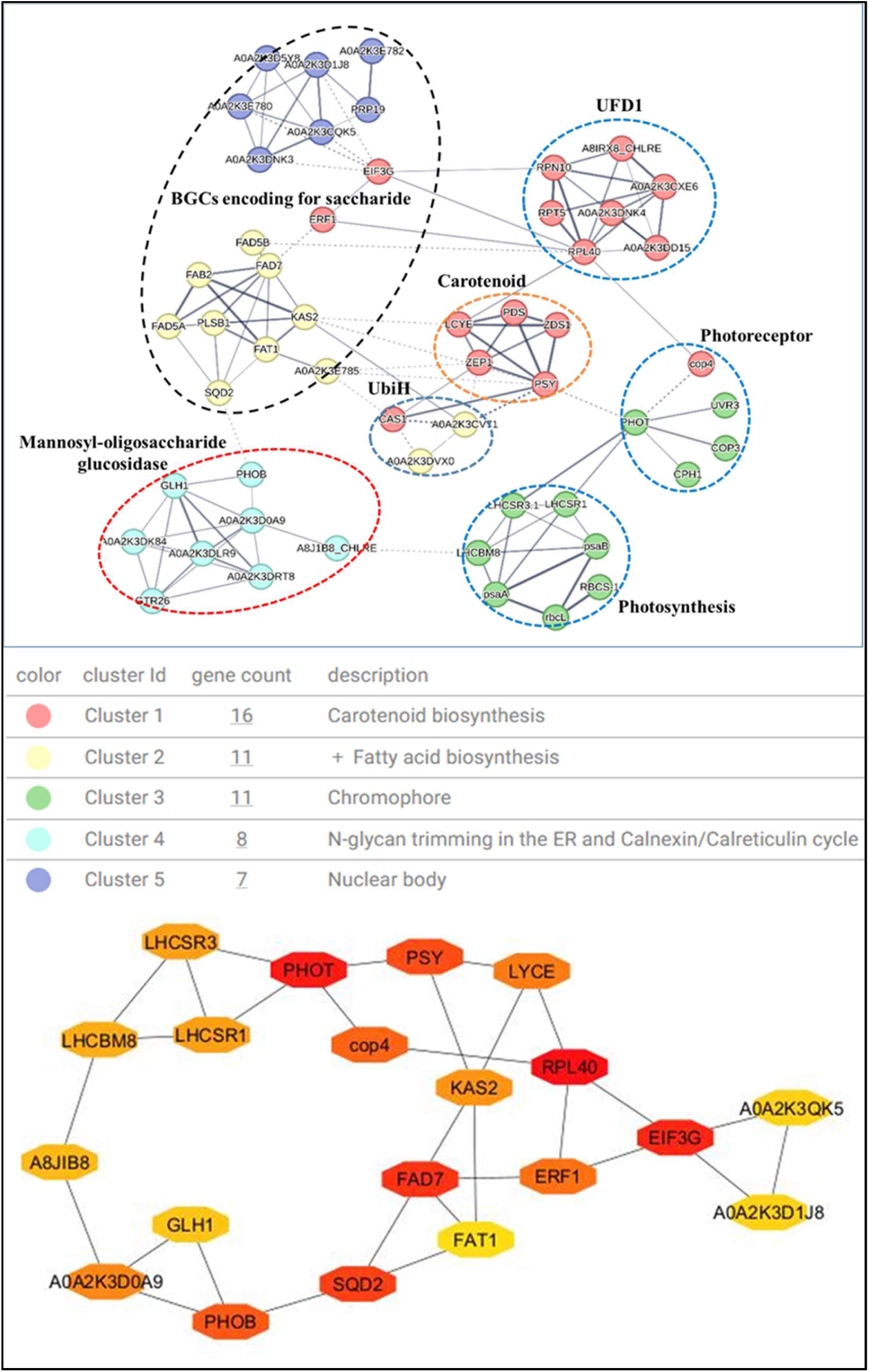
Represents bio-curated crosstalk between BGC encoding for saccharide and LOV-effector domains (mannosyl-oligosaccharide glucosidase, UFD1 and UbiH) The k-means clustering clustered the PPI into 5 clusters. The curated PPI was further visualised and analysed in Cytoscape (version 3.10.3) using the default layout with the top 20 betweenness methods and shortest path methods. All nodes that control the overall network were analysed using CytoHubba analysis using the betweenness algorithm. Colour range (red to yellowish) indicates scores ranked by the betweenness method with their interacting partners. PDS-Phytoene desaturase; Cop4- Chlamyopsin 4; PHOT- Phototropin, Blue light-sensing protein; FAD7-Chloroplast glycerolipid omega-3-fatty acid desaturase; ERF1-Eukaryotic release factor 1; EIF3G- Eukaryotic translation initiation factor 3 subunit G; PRP19- Spliceosome component, nuclear pre-mRNA splicing factor; SQD2- Uncharacterized protein, contains domain glycosyl_transferase_1-4; KAS2-3-oxoacyl-[acyl-carrier-protein] synthase; RPL40- Ribosome binding protein 40; LYCE- Lycopene ε-cyclase; A0A2K3D0A9-mannosyl oligosaccharide glucosidase; CAS- Terpene cyclase/mutase family member

**Fig. 6:**
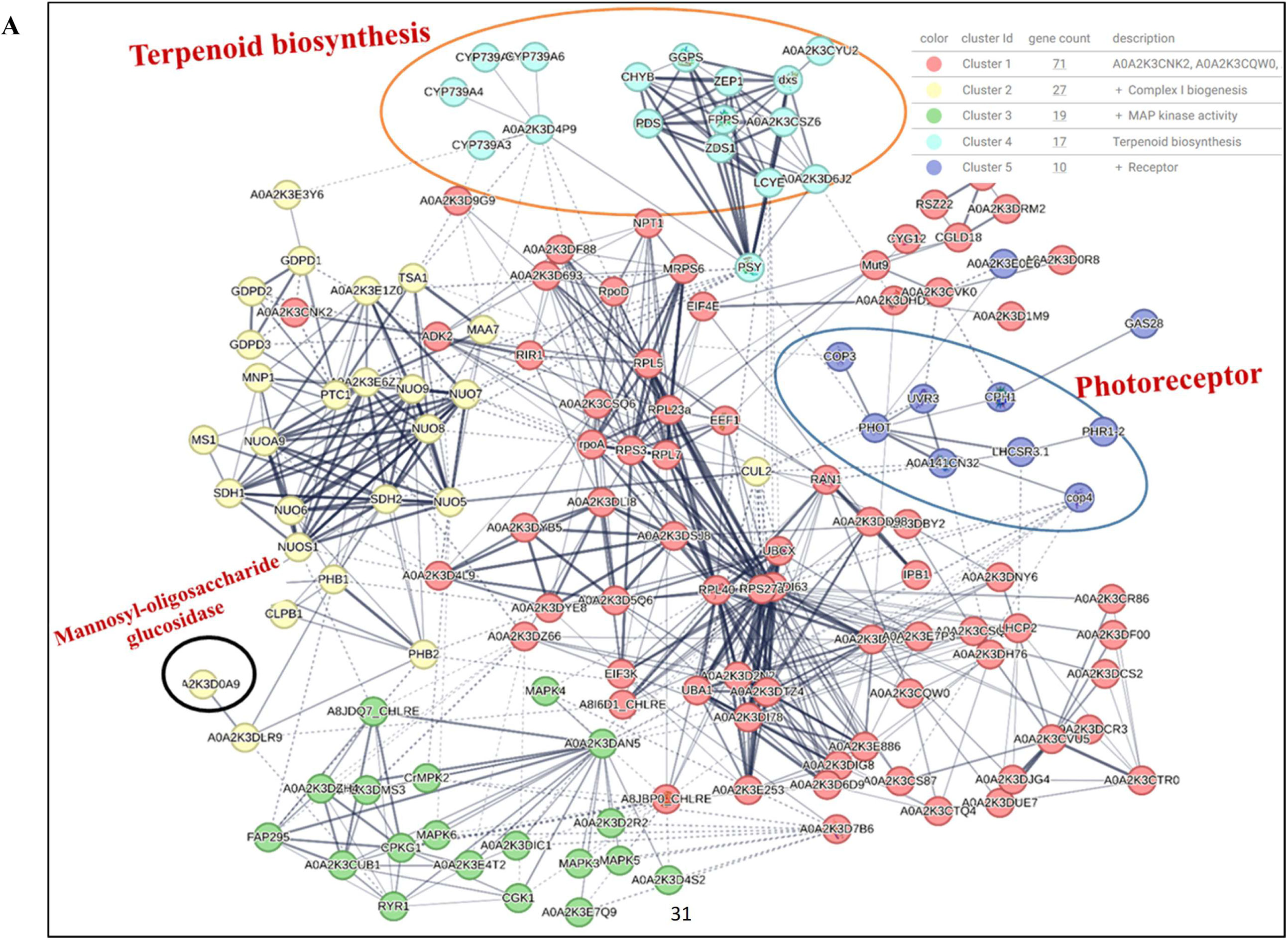
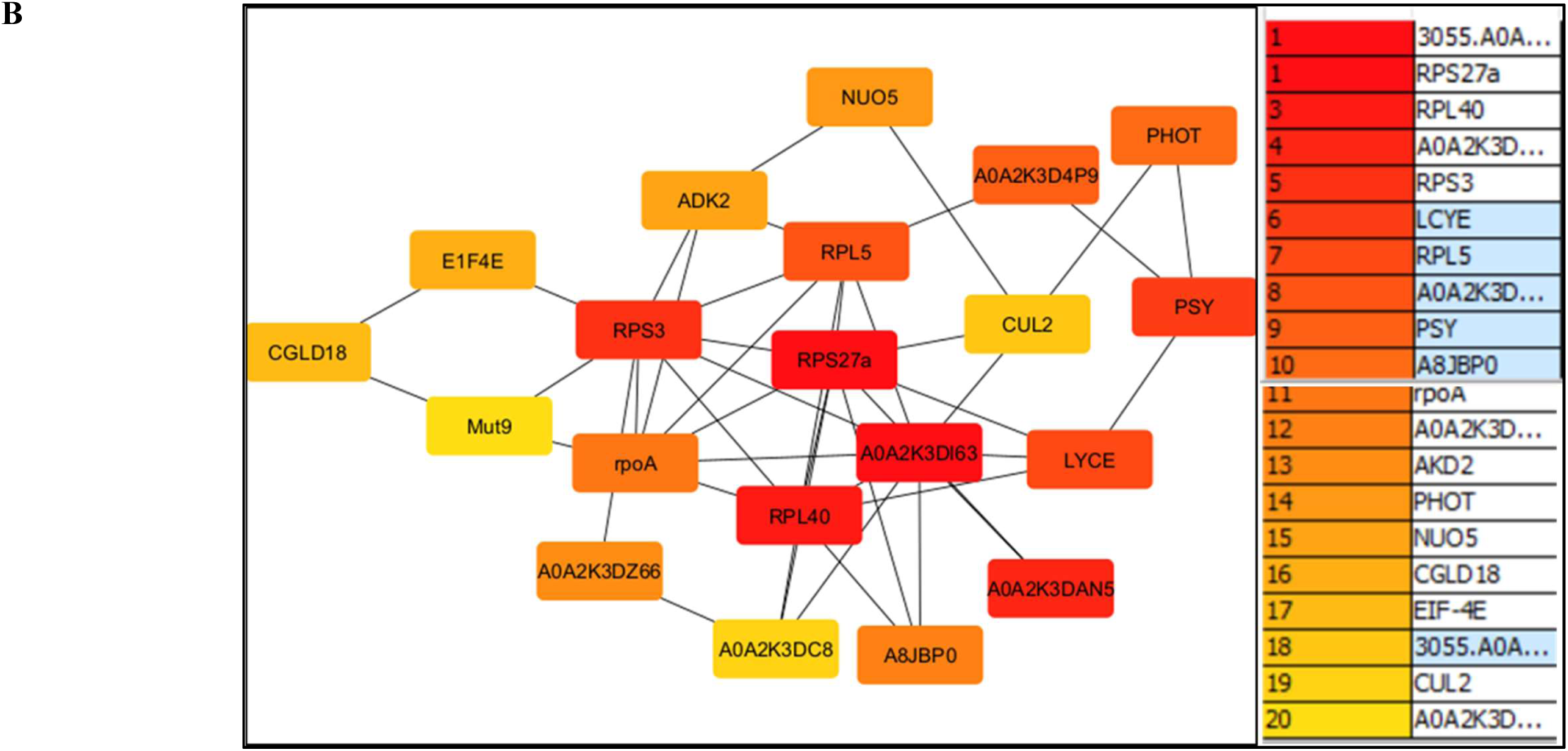
Bio-curated Photoreceptor, BGCs, and LOV-mannosyl oligoasaccharide glucosidase predicted protein networking in *C. reinhardtii*. (A) Shows crosstalk between BGC encoding for putative and LOV-mannosyl oligosaccharide glucosidase. The k-means clustering clustered the PPI into 5 clusters. (B) Shows all nodes that control the overall network were analysed using CytoHubba analysis using the betweenness algorithm and shortest pathway by applying default parameters. Colour range (red to yellowish) indicates scores ranked by the betweenness method with their interacting partners. Photoreceptor Cop4 shows a strong interaction via RPS27a, RPL40, A0A2K3DI63, A0A2k3DAN5 and PHOT with BGCs. These are also among the top 20 nodes affecting overall networking. Mannosyl oligosaccharide glucosidase also shows interaction with BGCs. However, it does not counts under the top 20 hub genes.

## 4 Conclusions and future perspectives

Our study unveils novel modular LOV-domain-containing proteins with the potential to function as light-inducible proteolytic, ribogenetic, and translational switches, enabling reversible and precise control over BGC-mediated metabolism by modulating illumination. The exploitation of the inherent modularity of LOV domains coupled with diverse effector domains provides a potential platform for light-controllable targeted cellular processes. These dynamic switches offer unprecedented spatiotemporal regulation of metabolic pathways. This innovation lays the foundation for next-generation opto-biotechnological tools, transforming light controlled synthetic biology, optogenetics and opto-biomanufacturing.

## Author Contributions

S.K. conceived the project. M. designed the experiments under the guidance of S.K. and performed the bioinformatics analysis. R.S. performed the biosynthetic gene cluster relevant analysis. R.S. drafted the original manuscript with the help of M. and S.K. All authors reviewed and approved the final manuscript.

## Author’s approval

All authors have seen and approved the manuscript. This work is original and not under consideration or published anywhere.

## Conflict of interest

The authors declare no competing interests.

## Funding sources

SK is thankful to ANRF/SERB, Government of India, for granting EEQ (EEQ/2023/000398) and CRG (CRG/2021/00315) research projects.

## Acknowledgements

SK is grateful to ANRF/SERB for providing research grants (EEQ/2023/000398 & CRG/2021/00315), respectively. RS acknowledge DBT-RA program (DBT-RA/2022/July/N/2560). M. acknowledge CSIR-UGC for providing the Junior research fellowship and Senior research fellowship (NTA Ref. No. 211610173246).

## Notes

### Competing Interest Statement

The authors have declared no competing interest.

