## Supplementary materials for "Novel algal modular LOV domain proteins expand opto-biotechnological avenues for controlling of eukaryotic riboswitching, translational and proteolytic processes"

**Supplementary Tables**

**Table S1: Important molecular determinants for chromophore binding and photocycle different LOV domains of modular proteins.**

| **CrLOV (Query)** | **N56** | **C57** | **R58** | **R63** | **Q61** | **N89** | **N99** | **Q120** |
| --- | --- | --- | --- | --- | --- | --- | --- | --- |
| LOV-IF4E | N | C | R | R | Q | N | N | Q |
| LOV-SppA | N | C | R | R | Q | N | N | Q |
| LOV-ABC transporter G-25 | N | C | R | R | Q | N | N | Q |
| LOV-PKsD | N | C | R | K | Q | N | N | Q |
| LOV-NTF2 | N | C | R | K | Q | N | N | Q |
| LOV-UFD1 | N | C | R | R | Q | N | N | G |
| LOV-HDAC Class II | N | C | R | R | Q | N | N | Q |
| LOV-UbiH | N | C | R | R | Q | N | N | Q |
| LOV-Mannosyl-oligossacharide glucosidase | N | C | R | R | Q | Q | N | Q |
| Lov-Peptidase S8_S53 | N | C | R | R | Q | N | N | Q |
| LOV-LrgB | N | C | R | R | Q | N | N | Q |
| LOV-XerD | N | C | R | R | Q | N | N | V |
| LOV-CifA/Voltage dependent potassium channel | N | C | R | Q | Q | N | N | Q |

#Red colour shows variation in amino acid residue

**Table S2: Protein Accession number and source organism used for homology analysis and phylogenetic analysis of different effector domains.**

| **eIEF4** | | | | | | | |
| --- | --- | --- | --- | --- | --- | --- | --- |
| Accession number | | Organism | | | | | |
| PTN18119.1 | | *Saccharomyces cerevisiae* | | | | | |
| XP_719387.1 | | *Candida albicans SC5314* | | | | | |
| XP_011779398.1 | | *Trypanosoma brucei gambiense DAL972* | | | | | |
| NP_648160.2 | | *Drosophila melanogaster* | | | | | |
| XP_018111588.1 | | *Xenopus laevis* | | | | | |
| NP_193538.1 | | *Arabidopsis thaliana* | | | | | |
| NP_001359122.1 | | *Oryza sativa Japonica Group* | | | | | |
| Pdb 5GW6 | | *Homo sapiens* | | | | | |
| GAQ81490.1 | | *Klebsormidium nitens* | | | | | |
| KAF5838073.1 | | *Dunaliella salina* | | | | | |
| **UbiH** | | | | | | | |
| Accession number | | | | Organism | | | |
| WP_007082050.1 | | | | *Rhodanobacter fulvus* | | | |
| WP_228326525.1 | | | | *Xanthomonas campestris* | | | |
| WP_126122971.1 | | | | *Pseudomonas (Multispecies)* | | | |
| MEH6452991.1 | | | | *Psychromonas sp. (MAG)* | | | |
| WP_027845304.1 | | | | *Mastigocoleus testarum* | | | |
| OIO57366.1 | | | | *Proteobacteria bacterium CG1_02_64_396 (MAG)* | | | |
| XP_042925688.1 | | | | *Chlamydomonas reinhardtii* | | | |
| GAQ86566.1 | | | | *Klebsormidium nitens* | | | |
| CAD5320460.1 | | | | *Arabidopsis thaliana* | | | |
| XP_015628965.1 | | | | *Oryza sativa Japonica Group* | | | |
| WP_136904379.1 | | | | *Rhodobacter capsulatus* | | | |
| EFK2607024.1 | | | | *Escherichia coli* | | | |
| **UFD1** | | | | | | | |
| Accession number | | | Organism | | | | |
| NP_001030324.2 | | | *Homo sapiens* | | | | |
| XP_006522069.1 | | | *Mus musculus* | | | | |
| pdb1ZC1 | | | *Saccharomyces cerevisiae* | | | | |
| XP_024387524.1 | | | *Physcomitrium patens* | | | | |
| NP_001031384.1 | | | *Arabidopsis thaliana* | | | | |
| NP_001396288.1 | | | *Oryza sativa Japonica Group* | | | | |
| RVW36573.1 | | | *Vitis vinifera* | | | | |
| GAQ88899.1 | | | *Klebsormidium nitens* | | | | |
| KAF5839060.1 | | | *Dunaliella salina* | | | | |
| XP_042916712.1 | | | *Chlamydomonas reinhardtii* | | | | |
| **Mannosyl oligosaccharide glucosidase** | | | | | | | |
| Accession number | | | | | Organism | | |
| XP_024341762.1 | | | | | *Postia placenta MAD-698-R-SB12* | | |
| PKK64666.1 | | | | | *Rhizophagus irregularis* | | |
| KAG1139740.1 | | | | | *Rhizopus arrhizus* | | |
| KAF6071613.1 | | | | | *Candida albicans* | | |
| KAF2733136.1 | | | | | *Polyplosphaeria fusca* | | |
| KAI8883093.1 | | | | | *Backusella circina FSU 941* | | |
| KAK0542742.1 | | | | | *Tilletia horrida* | | |
| XP_029737837.1 | | | | | *Sporisorium graminicola* | | |
| GFP65159.1 | | | | | *Saccharomyces cerevisiae* | | |
| AAC18786.1 | | | | | *Arabidopsis thaliana* | | |
| KAF2954024.1 | | | | | *Oryza sativa Japonica Group* | | |
| GAQ91059.1 | | | | | *Klebsormidium nitens* | | |
| KAF5826933.1 | | | | | *Dunaliella salina* | | |
| KAI2523967.1 | | | | | *Homo sapiens* | | |
| XBP80432.1 | | | | | *Escherichia coli* | | |
| **SppA, protease IV** | | | | | | | |
| Accession number | Organism | | | | | | |
| WP_353674719.1 | *Synechocystis sp. LKSZ1* | | | | | | |
| EHR8834757.1 | *Escherichia coli* | | | | | | |
| ELH8887442.1 | *Vibrio cholerae* | | | | | | |
| WP_105898824.1 | *Haemophilus influenzae* | | | | | | |
| XP_015626919.1 | *Oryza sativa Japonica Group* | | | | | | |
| NP_565077.2 | *Arabidopsis thaliana* | | | | | | |
| XP_042920246.1 | *Chlamydomonas reinhardtii* | | | | | | |
| WP_023244287.1 | *Salmonella (Multispecies)* | | | | | | |
| GAQ82938.1 | *Klebsormidium nitens* | | | | | | |
| **Peptidases S8_S53** | | | | | | | |
| Accession number | | | | | | | Organism |
| UHH08225.1 | | | | | | | *Bacillus subtilis* |
| QHB15709.1 | | | | | | | *Virgibacillus natechei* |
| WP_115512431.1 | | | | | | | *Xanthomonas arboricola* |
| KGQ06216.1 | | | | | | | *Beauveria bassiana D1-5* |
| XP_054562142.1 | | | | | | | *Fusarium oxysporum Fo47* |
| GAQ88137.1 | | | | | | | *Klebsormidium nitens* |
| GJN84156.1 | | | | | | | *Purpureocillium lilacinum* |
| HFQ2352909.1 | | | | | | | *Pseudomonas aeruginosa* |
| WP_181709490.1 | | | | | | | *Burkholderia (Multispecies)* |
| CAA75805.1 | | | | | | | *Aspergillus fumigatus* |
| XP_047606253.1 | | | | | | | *Trichophyton rubrum CBS 118892* |
| ETR97180.1 | | | | | | | *Trichoderma reesei RUT C-30* |
| **HDAC classII** | | | | | | | |
| Accession number | | | | | | Organism | |
| pdb 2VQO | | | | | | *Homo sapiens* | |
| NP_001398533.1 | | | | | | *Mus musculus* | |
| XP_024843702.1 | | | | | | *Bos taurus* | |
| XP_046764820.1 | | | | | | *Gallus gallus* | |
| Pdb 6UO7 | | | | | | *Escherichia coli* | |
| NP_001162760.1 | | | | | | *Drosophila melanogaster* | |
| NP_001257279.1 | | | | | | *Caenorhabditis elegans* | |
| NP_001190583.1 | | | | | | *Arabidopsis thaliana* | |
| KAF5831193.1 | | | | | | *Dunaliella salina* | |
| XP_015647003.1 | | | | | | *Oryza sativa Japonica Group* | |
| GAQ90206.1 | | | | | | *Klebsormidium nitens* | |
| EGA73404.1 | | | | | | *Saccharomyces cerevisiae AWRI796* | |

**Table S3: Table list the the hub genes of the bio-curated PPI-network.**

| Hub Gene(s) | Biosynthetic nature |
| --- | --- |
| PHOT | Phototropin, Blue light-sensing protein |
| A0A2K3D0A9 | Mannosyl oligosaccharide glucosidase domain |
| PSY | Chloroplast phytoene synthase |
| FAD7 | Chloroplast glycerolipid omega-3-fatty acid desaturase |
| LHCBM8 | Chlorophyll a-b binding protein |
| SQD2 | Uncharacterized protein, contains domain glycosyl_transferase_1-4 |
| PHOB | Alpha-1,4 glucan phosphorylase |
| A8J1B8 | ARF-GAP |
| KAS2 | 3-oxoacyl-[acyl-carrier-protein] synthase |
| ERF1 | Eukaryotic release factor 1 |
| CAS1 | Terpene cyclase/mutase family member; Belongs to the terpene cyclase/mutase family. |
| RPN10 | VWFA domain-containing protein. |
| RPL40 | Ribosomal protein L40. |
| A0A2K3DI78 | UBIQUITIN_CONJUGAT_2 domain-containing protein |
| COP3 | Chlamyopsin 3 |
| RPL5 | Ribosomal protein L5 |
| CUL2 | CULLIN_2 domain-containing protein; Belongs to the cullin family |
| NUO5 | NADH: ubiquinone oxidoreductase 24 kDa subunit |
| RpoD | Chloroplast RNA polymerase sigma factor |
| CGLD18 | B-box zinc finger protein |
| A0A2K3DZ66 | GP-PDE domain-containing protein |
| A0A2K3DKC8 | MgtE_N domain-containing protein |
| A0A2K3DAN5 | Uncharacterized protein |
| A0A2K3DI63 | Ubiquitin-like domain-containing protein |

**Table S4: List of different nodes and their annotations for the interacting partners of effector domains.**

| Node | | Annotation |
| --- | --- | --- |
| **Eukaryotic IF4E effector domain** | | |
| EIF4E | | Eukaryotic initiation factor; Belongs to the eukaryotic initiation factor 4E family |
| A0A2K3E7A5 | | Polyadenylate-binding protein; Binds the poly(A) tail of mRNA. Belongs to the polyadenylate-binding protein type-1 family. |
| EIF5Bbp | | eIF5Bbm |
| A0A2K3CNL9 | | MI domain-containing protein. |
| A0A2K3DL52 | | MIF4G domain-containing protein. |
| EIF3B | | Eukaryotic translation initiation factor 3 subunit B |
| A0A2K3D4R1 | | MI domain-containing protein |
| A0A2K3DIH9 | | Uncharacterized protein. |
| A0A2K3DQS2 | | Uncharacterized protein |
| FKB12 | | Peptidylprolyl isomerase |
| A0A2K3DMV5 | | Uncharacterized protein |
| **Ubiquitin fusion degradation 1 (UFD1) effector domain** | | |
| R1FJB3_EMIHU | *Ubiquitin fusion degradation 1 protein.* | |
| R1FV81_EMIHU | *Ubiquitin like domain containing protein* | |
| R1BK24_EMIHU | ANK_REP_REGION domain-containing protein | |
| R1DAU1_EMIHU | UAS domain-containing protein | |
| R1DK00_EMIHU | OTU domain-containing protein | |
| R1EPL0_EMIHU | Uncharacterized protein. | |
| R1CSN2_EMIHU | AAA domain-containing protein; Belongs to the AAA ATPase family. | |
| R1F4L3_EMIHU | MPN domain-containing protein. | |
| R1BRH9_EMIHU | VWFA domain-containing protein | |
| R1DBX9_EMIHU | VWFA domain-containing protein | |
| A0A0D3I624 | WD_REPEATS_REGION domain-containing protein. | |
| **UbiH effector domain** | | |
| Esi_0212_0023 | Kynurenine 3-Monooxygenase(UbiH) | |
| Esi_0036_0051 | Tryptophan synthase (Alpha / beta chains) | |
| Esi_0060_0108 | Tryptophan synthase (Alpha / beta chains) | |
| STE1 | Sterol C-5 desaturase; Belongs to the sterol desaturase family | |
| Esi_0048_0104 | Fatty acid hydroxylase domain-containing protein; Belongs to the sterol desaturase family | |
| Esi_0638_0003 | Uncharacterized protein. | |
| Esi_0115_0055 | Cytochrome b5 heme-binding domain-containing protein | |
| Nia | NAD(P)H-Nitrate reductase | |
| Esi_0302_0008 | Similar to Cytochrome b5 reductase 4; Belongs to the cytochrome b5 family | |
| CAS | Cycloartenol synthase. | |
| CAS-2 | Cycloartenol synthase. | |
| **Mannosyl oligosaccharide glucosidase effector domain** | | |
| GCSI | Mannosyl-oligosaccharide glucosidase, family GH63. | |
| Esi_0130_0015 | Calnexin. | |
| OST48 | Dolichyl-diphosphooligosaccharide | |
| RPN1 | Dolichyl-diphosphooligosaccharide | |
| GanAB | Alpha-glucosidase II alpha subunit, family GH31 | |
| Esi_0174_0043 | Uncharacterized protein. | |
| Esi_0056_0076 | Uncharacterized protein | |
| RPN2 | Dolichyl-diphosphooligosaccharide | |
| Esi_0177_0001 | Uncharacterized protein | |
| STT3 | annotation not available | |
| Esi_0529_0017 | annotation not available | |

**Supplementary Figures**

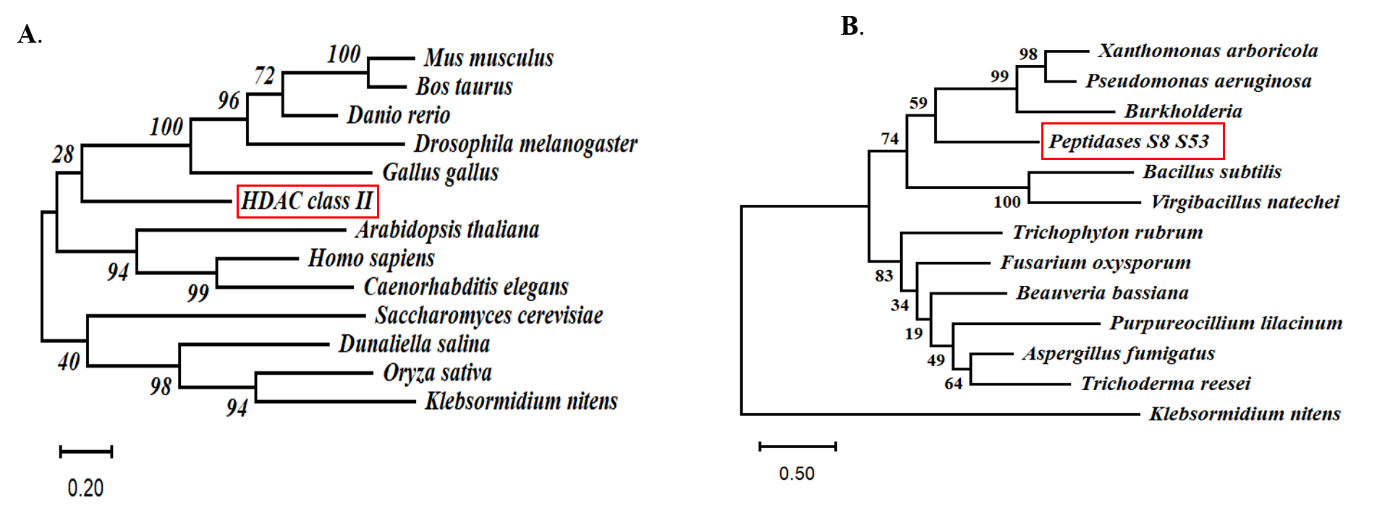

**Fig. S1:** Phylogenetic analysis of effector domain. (A) HDAC class II. (B) Peptidases S8 S53.

Alignment of protein sequences was carried out using ClustalW in MEGA12. The tree was analyzed using Maximum likelihood method and Jonas –Taylor-Thorton model with 1000 bootstrap replicates. Evolutionary analyses were conducted in using MEGA 12 software.

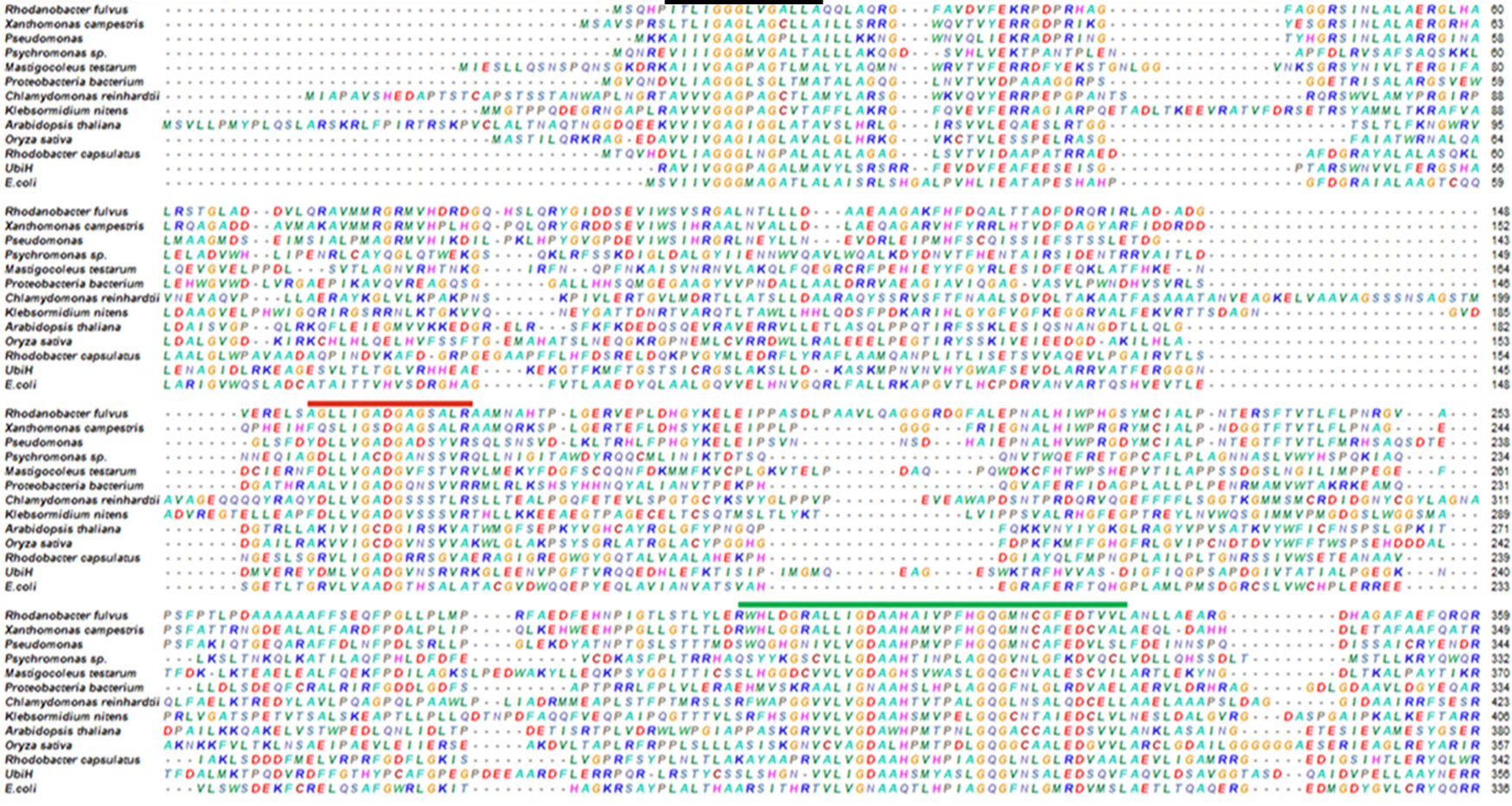

Fig. S2: - Homology analysis of UbiH effector domain. The black line indicate the first FAD (Flavin adenine dinucleotide) fingerprint sequence, which includes the well-known Rossmann fold (containing the GXGXXG motif), red line represents the DG amino acid sequence and green line represents the second FAD binding motif, containing the GD sequence with the highly conserved aspartyl residue that contacts the O-3′ of the ribose moiety of FAD.

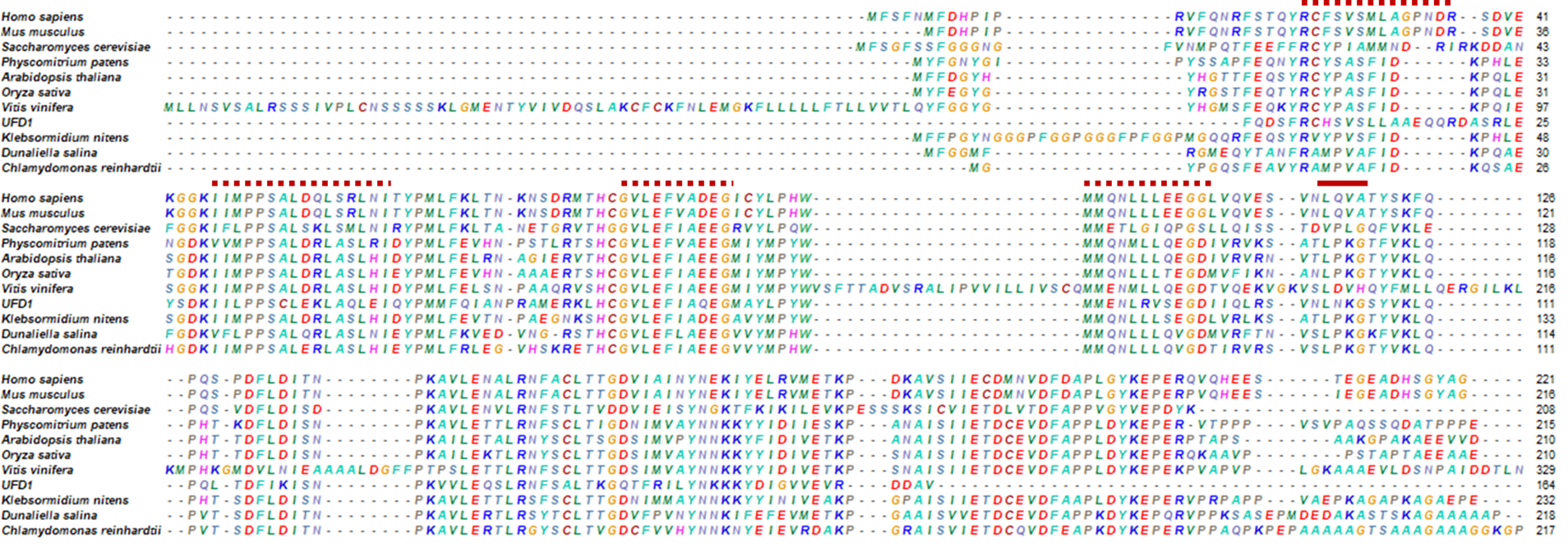

Fig. S3: Homology analysis for UFD1 effector domain. The red colour dashed line shows conserved double psi β-barrel motif. The red solid line represents conserved α/βroll motif.

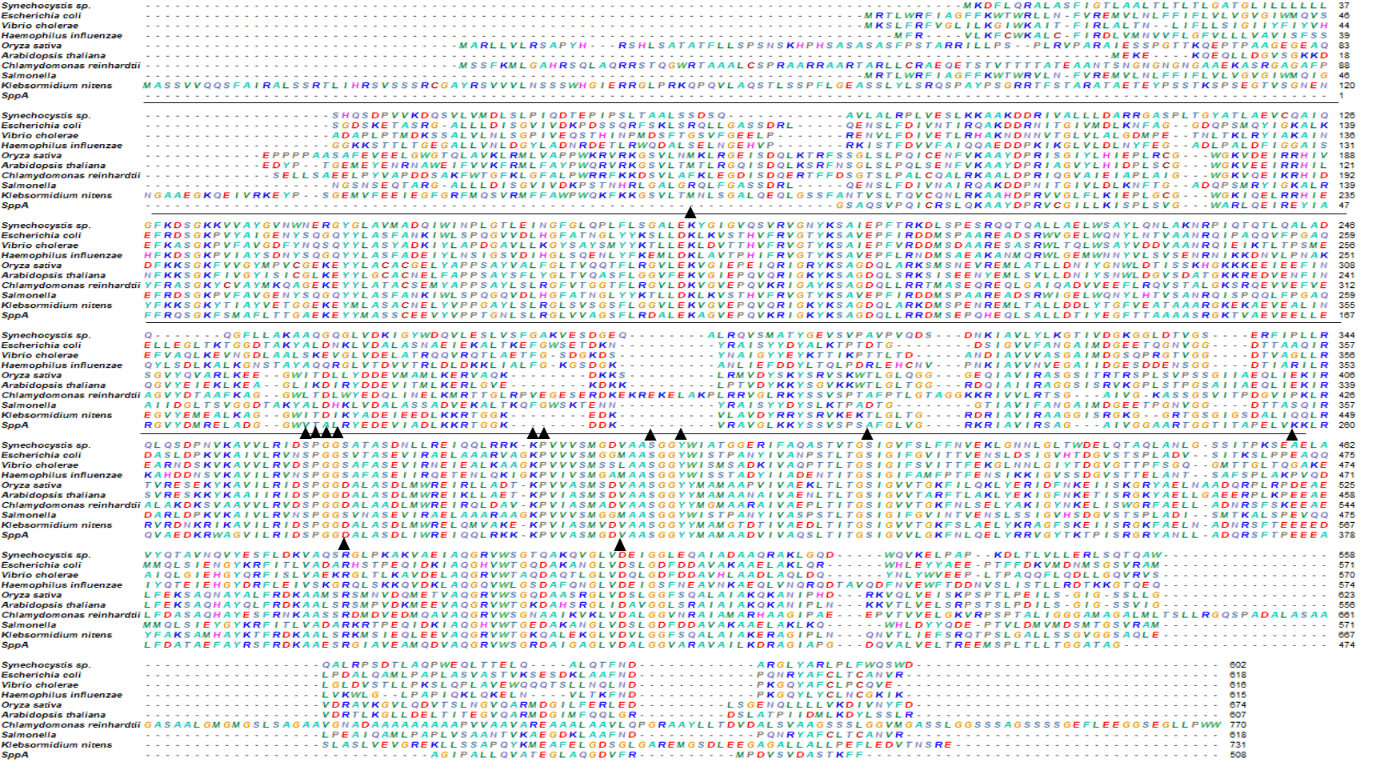
Fig. S4: Homology analysis of SppA, Protease IV. The triangle shows conserved residues that are present in all the sequences. The black line represents the amino terminal of the peptidase, and the remaining portion represents the carboxy terminus of the peptidase.

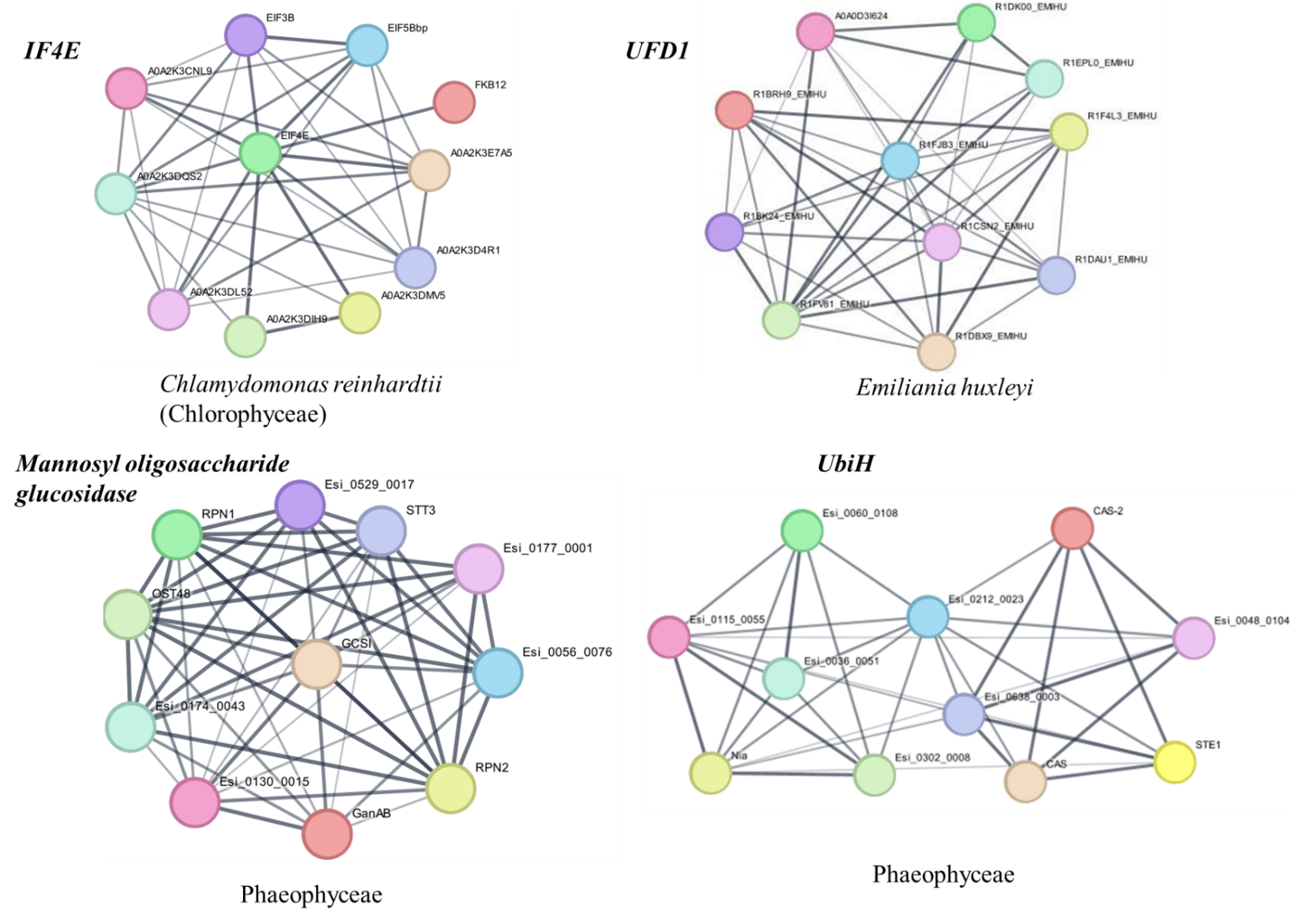

Fig. S5: Protein networking showing interacting partners of different effector domains. The interactome was predicted based on the closest lineage for the identified effector domains available in the String database.
